# EnvZ signal output correlates with capacity for transmembrane water encapsulation

**DOI:** 10.1101/2025.03.25.645265

**Authors:** Rahmi Yusuf, Joy Jun Yan Yau, George W. Baker-Valla, Robert J. Lawrence, Roger R. Draheim

## Abstract

In Gram-negative bacteria, porins span the outer membrane and control the influx of several prominent groups of antibiotics. Expression of these porins is often altered in clinical isolates exhibiting multidrug resistance. The major regulator of porin expression in *Escherichia coli* is EnvZ, a canonical sensor histidine kinase. It allosterically processes periplasmic interactions with MzrA and cytoplasmic osmosensing into a single unified change in the ratio of its kinase and phosphatase activities. To better understand how transmembrane communication is achieved by EnvZ, we began by performing disulfide crosslinking to monitor the position and dynamics of helices within the transmembrane domain in vivo and correlated these with modulation of signal output. Subsequently, we employed atomistic molecular dynamic simulations on full-length EnvZ embedded in a model Gram-negative membrane and observed that this in silico analysis accurately predicted the structure of the transmembrane domain embedded in the bacterial membrane. We conclude by proposing that the signal output of EnvZ correlates with its potential for water encapsulation within the transmembrane helical bundle and with unwinding of a helix adjacent to the periplasmic end of the second transmembrane helix. When combined with results from other bacterial receptors, it remains plausible that water encapsulation occurs in other topologically similar sensor kinases.

## INTRODUCTION

Recent modelling demonstrates that when antimicrobial resistance (AMR) remains left unchecked, 100 trillion USD worth of economic output will be lost and approximately 200 million people will expire prematurely by 2050 *(1, 2)*. Within this context of AMR, Gram-negative organisms are particularly burdensome as most antibacterial chemotherapeutics need to pass through a lipopolysaccharide (LPS)-coated outer membrane (OM) and those that gain entry are often pumped back out by efflux pumps *(3, 4)*. Thus, the OM serves as the first line of defence for Gram-negative bacteria as it is impermeable to large, charged molecules and influx is controlled by porins *(3, 5, 6)*.

Most porins involved in hydrophilic antibiotic transport by Gram-negative bacteria belong to the classical OmpF and OmpC families *(3)*. Transcription of these porins is governed by the intracellular concentration of phospho-OmpR (OmpR-P), which is controlled by EnvZ in response to changes in periplasmic interactions with MzrA and environmental osmolarity *(7– 11)*(Fig. S1A). At low intracellular levels of OmpR-P, transcription of *ompF* is upregulated, whereas at higher levels of OmpR-P, transcription of *ompF* is repressed and transcription of *ompC* is activated. This results in a predominance of OmpF at low osmolarity and OmpC at higher osmolarities or in the presence of MzrA *(12*–*14)* (Fig. S1B). Dramatic modification of porin balance, which has been observed within clinical isolates from patients undergoing treatment with antibacterials *(15*–*21)*, strongly supports further characterization of the underlying mechanisms of porin regulation by EnvZ. Furthermore, differential regulation of EnvZ has been shown to be involved in resistance to carbapenem *(22)*. Thus, in addition to maintenance of porin balance, EnvZ also plays other not well-understood roles in mediating antibacterial resistance that warrant further understanding.

From an extracellular perspective, MzrA, <u>m</u>odulator of Env<u>Z</u> and Omp<u>R</u> protein <u>A</u>, localises to the inner membrane and interacts with EnvZ within the periplasmic space *(11)*. It serves as an upstream regulator of EnvZ signal output that is independent of osmolarity, pH and procaine *(9)*. Within the cytoplasm, a fragment of EnvZ (EnvZ_c_), consisting of residues Arg^180^ through Gly^450^, has been shown to mediate physiologically appropriate responses to increasing NaCl and sucrose concentrations *in vitro* and to increasing sucrose *in vivo (23)*. These results demonstrate that the transmembrane domain (TMD) of EnvZ is responsible for allosteric coupling of sensory input from the attached periplasmic and cytoplasmic domains. Understanding how EnvZ transduces signal across the biological membrane would be a significant step toward understanding regulation of porin expression and direct manipulation of porin balance in Gram-negative bacterial cells.

To further understand the mechanism of transmembrane communication by EnvZ, we created a single-Cys-containing library of variants encompassing the TMD that connects the periplasmic and the cytoplasmic domains within EnvZ. We used *in vivo* disulfide crosslinking to monitor the position and dynamics of individual TM helices within stimulated and unstimulated EnvZ. The results demonstrate that our in silico system consisting of an EnvZ homodimer (EnvZ_2_) and a model Gram-negative inner membrane (G-IM) represents the homodimeric EnvZ TMD in native bacterial membranes. Analysis of this EnvZ_2_/G-IM system in conjunction with measuring signal output from single-Cys-containing mutants suggests that the transmembrane domain of EnvZ participates in water encapsulation to an extent that correlates with signal output. We also observed α-helical unwinding of residues adjacent to the periplasmic end of TM2 in silico and upon stimulus perception in vivo. We conclude by proposing that these water-coordinating motifs and helix-unwinding events might be conserved within topologically similar sensor histidine kinases.

## RESULTS

We previously created a version of EnvZ that had its sole Cys residue changed to an Ala residue (C277A) in order to assess the dynamics of the transmembrane domain (TMD) by disulphide crosslinking. We found that this Cys-less version of EnvZ possessed normal steady-state signal output and responded to environmental osmolarity in a similar manner to wild-type EnvZ. Using this Cys-less EnvZ as a template, we created a library of EnvZ variants that contained a single Cys residue from positions 11 to 41 and via disulphide crosslinking determined that no major rearrangements occurred along the TM1-TM1’ interface upon stimulus perception *(24)*. Residues Leu^160^ to Ile^181^ and Leu^160^ to Ile^179^ have been predicted by DGpred *(25)* and TMHMM v2.0 *(26)* to comprise TM2 respectively and here, based on these predictions, we expanded our library to include variants with a single Cys residue from positions 156 to 184 (Fig. S2A). We observed that these new variants were stable when expressed within *E. coli* cells grown either under the low- or high-osmolarity regimes (Fig. S2B).

### Loss of sulfhydryl-reactivity adjacent to the periplasmic end of TM2 upon stimulus perception

In a similar manner to mapping TM1-TM1’ interactions*(24)*, Cys-containing EnvZ variants were expressed in *E. coli* cells and upon entering the early exponential phase, were subjected to 250 μM molecular iodine for 10 minutes and analyzed by non-reducing SDS-PAGE and immunoblotting (Fig. 1A). Within TM2, three distinct regions were observed. The N-terminal region (region I in Fig. 1B), comprised of residues 156 to 161, exhibited extensive cross-linking under the low-osmolarity regime (0% sucrose) and minimal crosslinking under the high-osmolarity (15% sucrose) regime. It should be noted that region I should be considered periplasmic because residue position 161 appears to be the final periplasmic position as a Cys residue at this position appears to dimerize in absence of additional oxidizing agent (Fig. S2B). The second region (II) consisting of positions 162 to 179, demonstrated reduced crosslinking near the boundaries and enhanced crosslinking within the membrane core, in a pattern similar to TM1-TM1’ reactivity *(24)*. The final region (III), from residues 180 to 184, showed no crosslinking (Fig. 1B). This significant osmolarity-mediated change, the loss of crosslinking at the at the periplasmic end of TM2 (red dashed box in Fig. 1B) was not observed during similar analyses of the TM1-TM1 interface and represents the sole osmolarity-mediated difference in cross-linking observed *(24)*.

**Fig 1.**
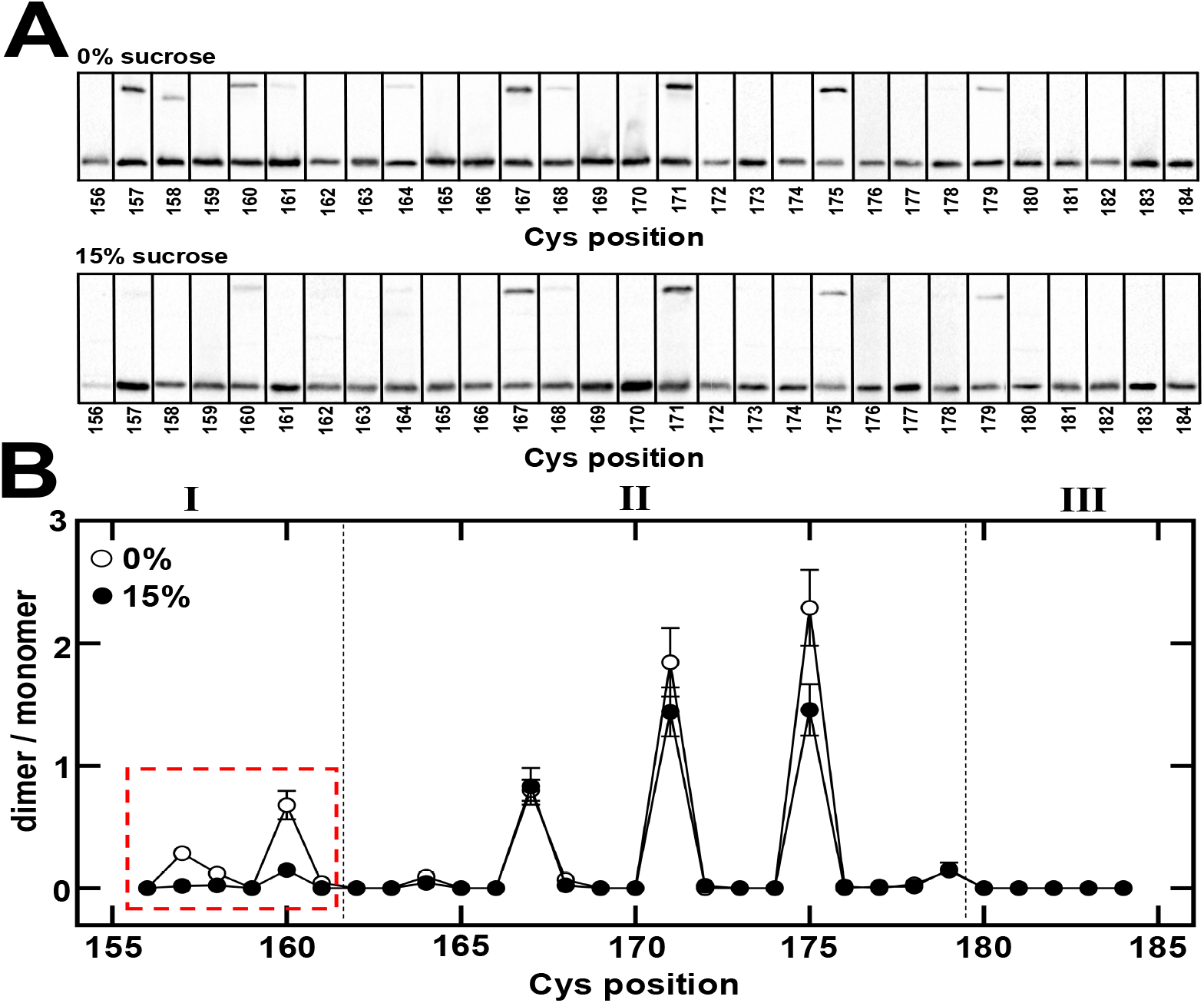
Mapping TM2-TM2’ interactions by sulfhydryl-reactivity analysis. (A) EPB30/pRD400 cells expressing one of the single-Cys-containing EnvZ receptors were grown under the low- (0%) or high-(15%) regimes until an OD_600nm_ of approximately 0.25 was reached. Cultures were then subjected to 250 μM molecular iodine. Particular Cys-containing EnvZ receptors resulted in the presence of dimeric EnvZ moieties that migrated at a slower rate than the monomeric species. (B) A minimum of three immunoblots were used for each of the data points present this allowed us to determine the dimer/monomer ratio represented on the Y-axis. Three distinct regions, denoted I, II and III were observed and are described in the text. Error bars represent the standard error of the mean with a sample size of n ≥ 3. The loss of α-helicity upon stimulus perception (15% sucrose) is denoted by a red dashed box.

### EnvZ signal output modulated based on Cys position

To assess signal output from the single-Cys-containing variants, we began by expressing them in EPB30/pRD400 cells, which allowed us to measure CFP fluorescence, YFP fluorescence, and to calculate the CFP/YFP ratio that estimates steady-state EnvZ signal output (Fig. S1B). Cells expressing the Cys-less EnvZ C277A were used as a baseline comparison as described previously *(24)*. When these cells are grown under the low-osmolarity regime, a shift in signal output toward the “on”, or kinase-dominant state, results in increased CFP fluorescence, reduced YFP fluorescence and an increase in the overall CFP/YFP ratio, while a shift toward the “off”, or phosphatase-dominant state, appears as decreased CFP, increased YFP and a decrease in CFP/YFP ratio (Fig. S1B).

When cells were grown under the low-osmolarity regime, EnvZ was less tolerant of Cys substitutions at the N- and C-terminal regions of TM2. Signal output from receptors containing a Cys at positions 156, 162 and 163 were elevated, exhibiting greater than a 5-fold increase in CFP/YFP, while receptors possessing a Cys at cytoplasmic positions 181, 182 and 184 possessed over a 2-fold increase in CFP/YFP. These boundary regions appear to flank a core of alternating increases and decreases in EnvZ signal output, as observed between residue positions 165 and 180 (Figs. 2A and 2B). When grown under the high-osmolarity regime, a pattern appeared where the Cys substitutions resulted in significant decreases in signal output (Fig. 2C). Of the 29 mutants analyzed, nearly half supported less than 75% of the wild-type signal output when grown under the high-osmolarity regime.

**Fig 2.**
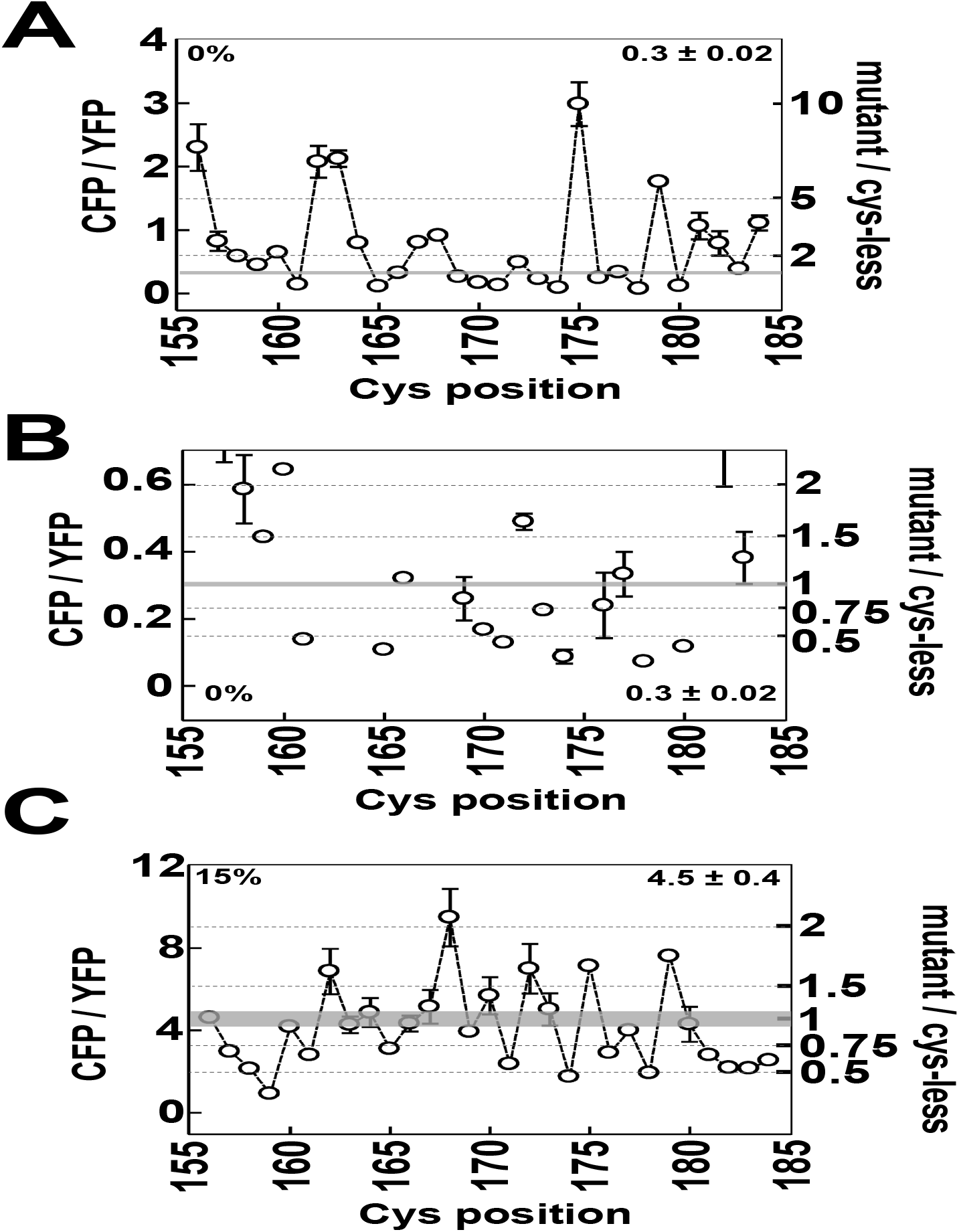
Signal output from the library of single-Cys EnvZ variants. (A) CFP/YFP from EPB30/pRD400 cells expressing one of single-Cys variants grown under the low-osmolarity (0% sucrose) regime. On the right axis, these CFP/YFP ratios are compared to the Cys-less (C277A) variant. (B) Magnified version of panel A in order to emphasise the region up to a 2-fold increase in CFP/YFP. (C) CFP/YFP from EPB30/pRD400 cells expressing one of the single-Cys variants grown under the high-osmolarity (15% sucrose) regime. On the right axis, these CFP/YFP ratios are compared to the Cys-less (C277A) variant. The shaded areas represent the mean signal output from the Cys-less variant of EnvZ with a range of one standard error of the mean. These values are provided to aid in comparison. Error bars represent standard deviation of the mean with a sample size of n ≥ 3.

### The EnvZ_2_/G-IM system accurately represents in vivo structure of the TMD

Due to the complex nature of these results, the differences of the TM1-TM1’ and TM2-TM2’ interfaces and the difficulty involved in comparing both in parallel, we turned to structural prediction and downstream atomistic molecular simulation. We began by subjecting the wild-type sequence of EnvZ from *E. coli* K-12 MG1655 to AlphaFold-Multimer v2.3.0 *(27, 28)* in order to produce three-dimensional coordinates of a homodimeric EnvZ, hereafter referred to as EnvZ_2_. A protein/membrane system (Fig. S3) composed from a single EnvZ_2_ moiety and a model Gram-negative inner membrane (G-IM) *(29*–*32)* was created with Membrane Builder *(33*–*35)*, subjected to the default six-step equilibration protocol with NAMD 2.14, and NAMD 3 was employed for all subsequent simulation *(36)*.

We assessed the quality of the TMD generated by comparing the inter-residue distances from the EnvZ_2_/G-IM system to our previous TM1-TM1’ crosslinking data *(24)* and the TM2-TM2’ results generated here (Fig. 3). The extent of *in vivo* crosslinking is based upon the position and orientation of the Cys side chains, and we predicted an inverse relationship between the extent of crosslinking and distance between the reacting sulfhydryl groups. Residues within TM1 and TM2 that form local distance minima were found to represent localized crosslinking maxima, including residues positions 11, 15, 19, 23, 26, 30 and 34 along the TM1-TM1 interface and residue positions 157, 160, 164, 167, 171, 175 and 179 along the TM2-TM2 interface (red lines in Fig. 3). Within the EnvZ_2_/G-IM system, these residue positions face inward toward the core of an antiparallel four-helix bundle that forms the TMD of EnvZ_2_ (Fig. S4). Overall, these results demonstrate that the EnvZ_2_/G-IM system represents the homodimeric EnvZ TMD in native bacterial membranes.

**Fig 3.**
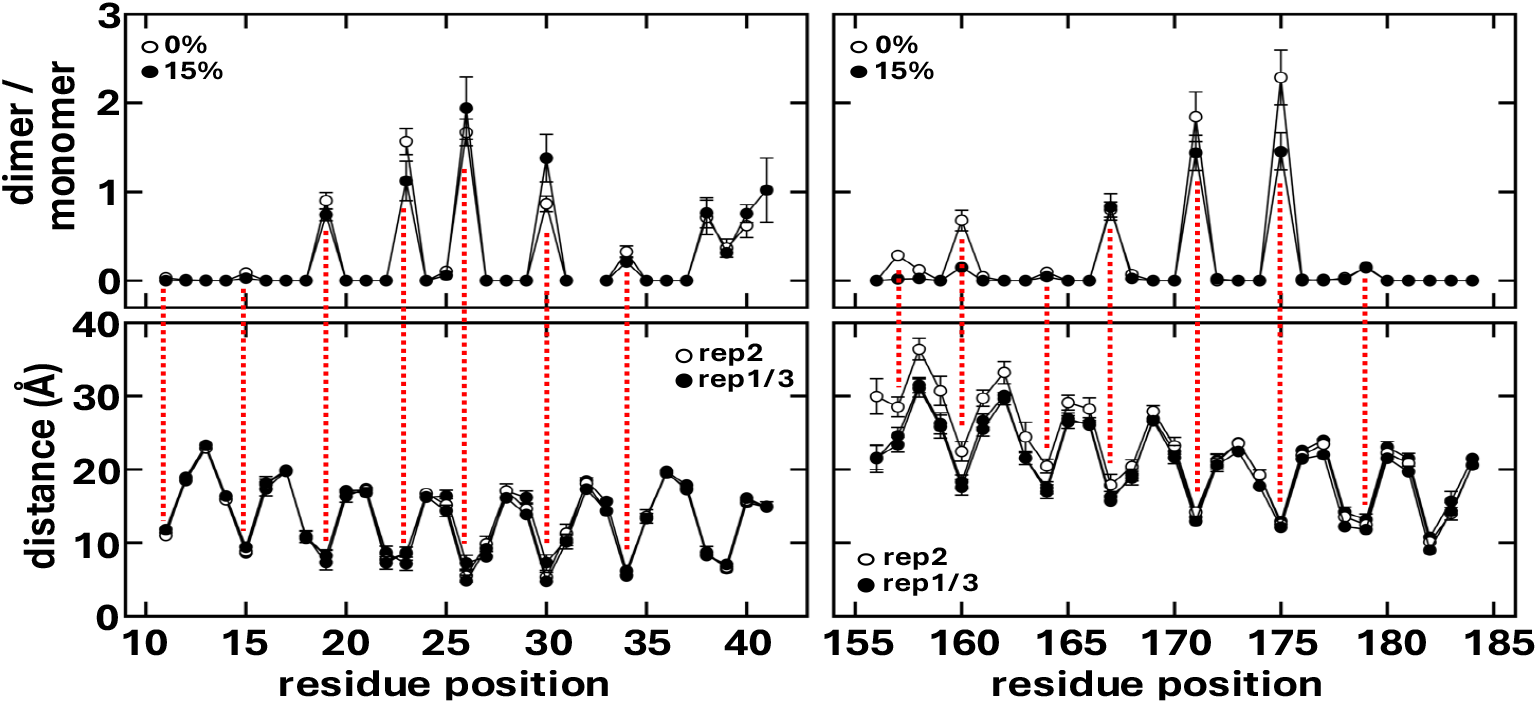
Comparison of inter-residue distances by sulfhydryl-reactivity analysis in vivo and within the three replicates of the EnvZ_2_/G-IM simulation. (top) Dimer-monomer ratio estimate inter-residue distances between the side chains within an EnvZ homodimer, i.e. Cys11 and Cys11’. The TM1-TM1’ data was previously published *(24)*, while the TM2-TM2’ data is shown in Fig. 1B. (bottom) The average interresidue distance between the same side chains within the EnvZ homodimer are shown from three replicate simulations of the same EnvZ_2_/G-IM system. These orthogonal methods show (red dashed lines) that localized crosslinking maxima correlate with the localized distance minima in the simulations.

## DISCUSSION

### Previous evidence for water coordination within the TMD of SHKs

Several previous studies support water encapsulation within the TMD of SHKs. The earliest support comes from the *E. coli* PhoQ sensor kinase that was shown to require a hydrophilic residue within its TMD *(37)*. Interpretation within the context of the previously identified structure of HtrII from *Natronomonas pharanois*, which possesses a cytoplasmic-facing water-filled hemichannel *(38)*, with subsequent support from molecular simulations with an assembled PhoQ TMD *(39)*, led to the proposal that PhoQ possesses a cytoplasmic-facing water-filled hemi-channel within its TMD. Lending credence to this proposal was the high-resolution crystal structure of TMD-containing *E. coli* NarQ receptor, which was shown to possesses water within its TMD, supporting the idea that this membrane-embedded hemi-channel is not solely found in the PhoQ and HtrII TMDs *(40)*. A preprint examining atomistic MD simulations of a membrane-embedded *E. coli* PhoQ has shown a high level of hydration within the TMD and it has been suggested that this represents an intermediate signaling state *(41)*. In summary, these previous results support the presence of a cytoplasmic-facing hemi-channel in several topologically similar bacterial proteins.

### Capacity for water encapsulation correlates with EnvZ signal output

To assess the appropriateness of water encapsulation within EnvZ, we examined signal output from our library of single-Cys-containing EnvZ variants within its context. To complete this analysis topologically, from the cytoplasm to the periplasm, we combined the data derived from our previous analysis of TM1 *(24)* and the data produced here covering TM2. As shown in Figure 4A, we have highlighted positions where a Cys residue results in greater than double the baseline EnvZ signal output, as indicated by a CFP/YFP ratio of greater than 0.6 (shaded in green in Fig. 4A). From this data, three motifs have been shown to be important for modulation of EnvZ signal output. The first is the cytoplasmic-facing water-filled hemichannel where a depth-based Gaussian of signal outputs is observed for Cys residues scanned from position 11-18. Two exceptions from this range, namely positions 12 and 17, possess side chains that do not face inward toward the solvent-filled hemi-channel. When a Cys residue is scanned along TM2, the side chains from positions 175 and 179 protrude into the hemi-channel and thus also stimulate EnvZ signal output. These residue positions are highlighted in green in Fig. 4A. To assess how far this cytoplasmic-facing water-filled hemichannel might protrude into the cytoplasm, we scanned Cys residues from positions 2 to 10. No cross-linking was observed when EPB30/pRD400 cells were grown under the low- or high-osmolarity regimes (Fig. S5). A large increase in signal output was only observed at position 10 both under the low-and high-osmolarity regime (Fig. S6). The water encapsulation motif is composed of two sets of Ser^26^, Thr^30^ and Thr^164^ residues (cyan residues in the middle panel of Fig. 4A). Significant increases in EnvZ signal output were observed at positions where the Cys residue would expand the water encapsulation pockets by facing toward the same bundle core but one helical turn lower, namely position 23 in TM1 or positions 167 and 168 in TM2. These positions have been highlighted in green in Fig. 5 Residue position 27 would also expand the water encapsulation pocket and is highlighted in green. The final motif consists of the periplasmic face of the TMD where Cys residues might be found within the periplasm. We have estimated where this might occur based on crosslinking observed in the absence of oxidizing agent (dashed lines in the top panel of Fig. 4). Thus, it appears that EnvZ is activated in when a Cys is found at positions 162 and 163, which face outward from the bundle and it is possible that a Cys residue here is facilitating water entry into the encapsulation motifs. These are the only two residue positions facing outward where a Cys substitution appears to simulate EnvZ. In contrast, we have also mapped the positions where Cys substitution inhibit EnvZ output below 75% wild-type output (Fig. 4B). In all cases, these residue positions reducing signal output fail to face into the helical core of EnvZ TMD.

**Fig 4.**
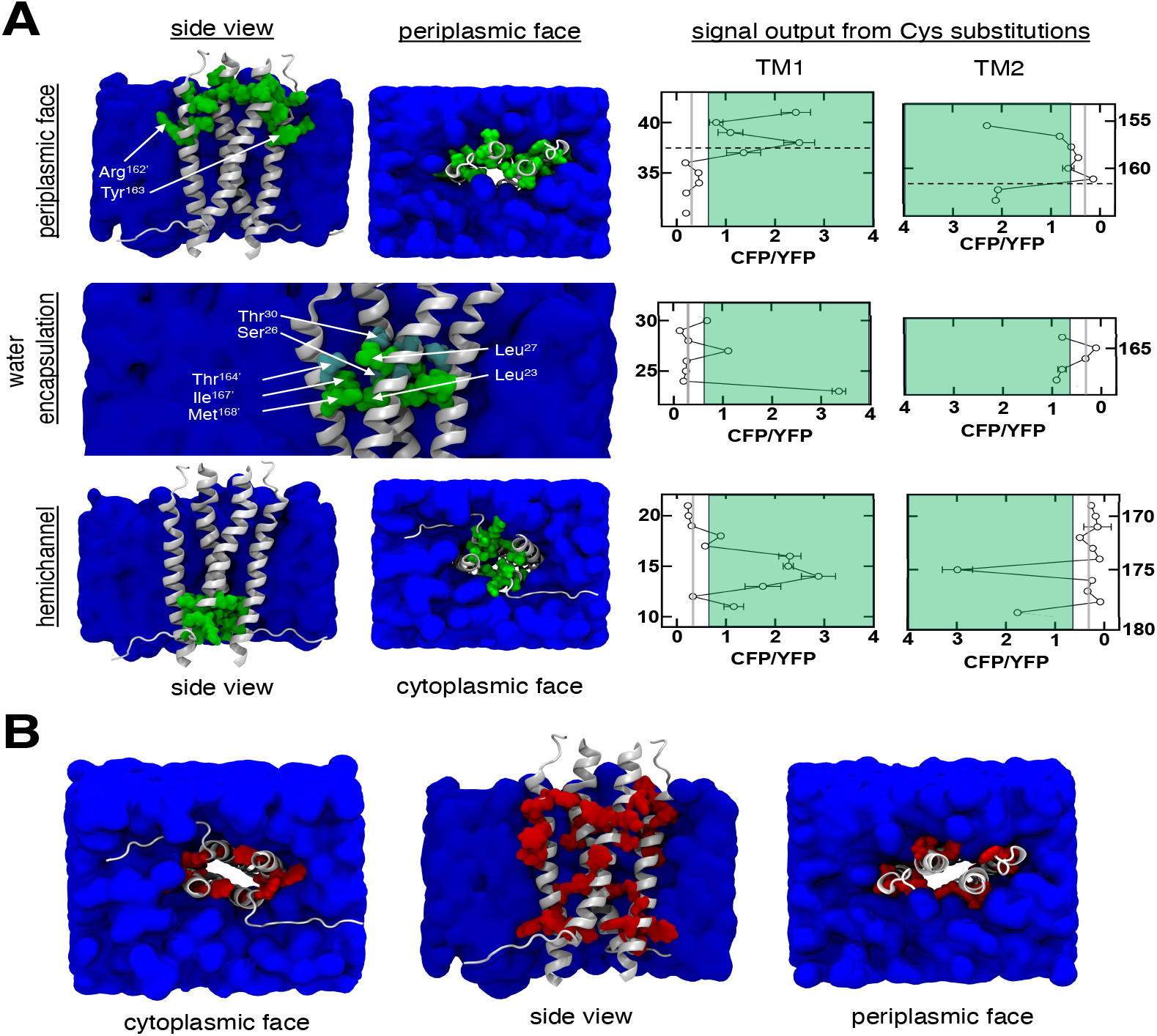
Facilitation of water encapsulation governs EnvZ signal output. (A) Alterations of the three water-coordination motifs modulate signal output. Residue positions where Cys substitutions activate signal output two-fold above baseline (CFP/YFP > 0.6) are highlighted in green. Three motifs have been identified where this occurs: when the Cys residues face into the cytoplasmic-facing water-filled hemichannel (lower third), when the water-encapsulation pockets are expanded (middle third) or when one of two outward-facing members of the periplasmic anchoring belt are substituted (upper third). (B) Residue positions when a Cys substitution reduces signal output to below 75% of baseline output are highlighted in red. Please note that none of these sidechains face into the transmembrane bundle core.

### Tuning signal output with strategically positioned Cys residues

We, and many other research groups, have attempted to modulate signal output from receptors in a broadly applicable and easy-to-use manner. In our case, we looked at the role of amphipathic aromatic residues and their preference for the water-hydrophobic interfaces of lipid bilayers, an approach that we referred to as “aromatic tuning.” According to this approach, amphipathic aromatic residues, namely Trp and Tyr, can be used to physically reposition the end of the TM2 with respect to the membrane. We showed that this method functioned in a linear manner within the aspartate chemoreceptor of *E. coli*, but not in the case of EnvZ *(42*–*49)*. Thus, aromatic tuning was not as widely applicable across topologically similar receptors as we initially sought. Here, we show that a single Cys residue in the cytoplasmic tail of EnvZ modulates signal output in a predictable manner suggesting that the cytoplasmic-facing, water-filled hemichannel might be used to activate signal output from any bacterial receptor of interest that shared membrane-spanning topology with EnvZ.

## MATERIALS AND METHODS

### Bacterial strains and plasmids

*E. coli* strains DH10B (New England Biolabs) or MC1061 *(50)* were used for DNA manipulations, while strain K-12 MG1655 *(51)* served a non-fluorescent strain that was used to control for light scattering and cellular autofluorescence. *E. coli* strains MDG147 [MG1655 Φ*(ompF*^*+*^-*yfp*^*+*^) Φ*(ompC*^*+*^-*cfp*^*+*^)] *(52)* and EPB30 (MDG147 *envZ::kan) (53)* were employed for analysis of EnvZ signal output. As the C-terminus of bacterial receptors can be sensitive to the presence of an epitope tag, we previously ensured that the addition of a V5-epitope tag did not alter the signaling properties of EnvZ *(43, 54)*. Plasmid pEB5 was employed as an empty control vector *(55)*. Plasmid pRD400 *(43)* retains the IPTG-based induction of EnvZ from plasmid pEnvZ *(56)* while adding a seven-residue linker (GGSSAAG) *(57)* and a C-terminal V5 epitope tag (GKPIPNPLLGLDST) *(58)*.

### Analysis of sulfhydryl-reactivity in vivo

Bacterial cultures were grown as described previously *(43)* with minor modification. MDG147 or EPB30 cells were transformed with pRD400 expressing one of the single-Cys-containing EnvZ receptors or pEB5 (empty). Fresh colonies were used to inoculate 2-ml overnight cultures of minimal medium A *(59)* supplemented with 0.2% glucose. Ampicillin, sucrose and IPTG were added as appropriate. Cells were grown overnight at 37 °C and diluted at least 1:1000 into 7 ml of fresh medium. Upon reaching an OD_600nm_ ≈ 0.3, cells were subjected to between 250 μM molecular iodine for 10 min while incubating at 37 °C. The reaction was terminated with 8 mM N-ethylmaleimide (NEM) and 10 mM EDTA. Cells were harvested by centrifugation and resuspended in standard 6X non-reducing SDS-PAGE buffer supplemented with 12.5 mM NEM. Cell pellets were analysed on 10% SDS/acrylamide gels. Standard buffers and conditions were used for electrophoresis, immunoblotting and detection with enhanced chemiluminescence *(60)*. Anti-V5 (Invitrogen) was the primary antibody and peroxidase-conjugated anti-mouse IgG (Sigma) was the secondary antibody. Digitized images were acquired with a ChemiDoc MP workstation (Bio-RAD), analysed with ImageJ v1.54j *(61)* and quantified with QtiPlot v1.2.2.

### Analysis of EnvZ signal output in vivo

Cells were grown as above with a few slight differences. Upon reaching an OD_600nm_ ≈ 0.3, chloramphenicol was added to a final concentration of 170 μg/ml. Fluorescent analysis for datasets assessing TM1 and TM2 were immediately conducted with 2 ml of culture and a Varian Cary Eclipse (Palo Alto, CA). For analysis of the library where the Cys residue was placed between residues positions 2 and 10, 5 ml of culture were centrifuged and resuspended into a 96-well plate for analysis within a Varian Cary Eclipse (Palo Alto, CA). CFP fluorescence was measured using an excitation wavelength of 434 nm and an emission wavelength of 477 nm, while YFP fluorescence was measured using an excitation wavelength of 505 nm and an emission wavelength of 527 nm. These values were corrected for cell density and for light scattering/cellular autofluorescence by subtracting the CFP and YFP fluorescence intensities determined for MG1655/pEB5 cells.

### Atomistic MD simulations and analysis

The wild-type sequence of EnvZ from *E. coli* K-12 MG1655 was submitted to AlphaFold-Multimer v2.3.0 *(27, 28)* and the output served as a homodimeric EnvZ (EnvZ_2_) moiety to embed within a model Gram-negative inner membrane *(29*–*32)*. This EnvZ_2_/G-IM system was generated by Membrane Builder and subsequent NAMD input files were generated by CHARMM-GUI *(33– 35)*. The initial system size and composition is presented in Fig S3. The default six-step equilibration protocol from Membrane Builder was employed. Production runs were performed for a minimum of 200 ns using a 2 fs timestep and in the absence of hydrogen mass partitioning. van der Waals interactions were cutoff at 12 Å with a force-switching function between 10 and 12 Å *(62)* and electrostatic interactions were calculated by the particle-mesh Ewald method *(63)*. The temperature was assigned at 303.15 K and the pressure (at 1 atm) were controlled by Langevin dynamics with a friction coefficient of 1 ps^−1^ *(64)*. MDAnalysis was used for downstream analysis *(65)*.

## Supplementary Materials

Fig. S1. Monitoring modulation of EnvZ signal output upon stimulus perception.

Fig. S2. Creation of a new library of single-Cys-variants that encompasses TM2.

Fig. S3. Composition and dimensions of the EnvZ_2_/G-IM system.

Fig. S4. Mapping the TM1-TM1’ and the TM2-TM2’ interface within the EnvZ TMD.

Fig. S5. Absence of cross-linking from the cytoplasmic tail single-Cys-variants.

Fig. S6. Signal output from EnvZ receptors containing a Cys residue in the cytoplasmic tail.

## Acknowledgments and Notes

We acknowledge suggestions on improvement of the manuscript from Christian Jørgensen (University of Portsmouth) and C. Keith Cassidy (University of Missouri). Robert Lawrence acknowledges support in the form of a doctoral bursary while Roger Draheim was supported with start-up funding from the Faculty of Science and Health and from the Institute of Biological and Biomedical Sciences (IBBS) at the University of Portsmouth.

## Author contributions

R.R.D. designed the project. R.Y., G.W.B. and R.R.D. were responsible for design of the methodology. G.W.B and R.R.D. were responsible for the software employed during the project. R.Y., J.J.Y.Y., G.W.B., R.J.L. and R.R.D. carried out the investigation and formal analysis. R.Y., R.J.L. and R.R.D. provided resources and acquired funding. The original draft was written by R.Y. and R.R.D, while review and editing was conducted by R.Y., J.J.Y.Y., G.M.B., R.J.L. and R.R.D.

## Supplementary material

**Figure S1.**
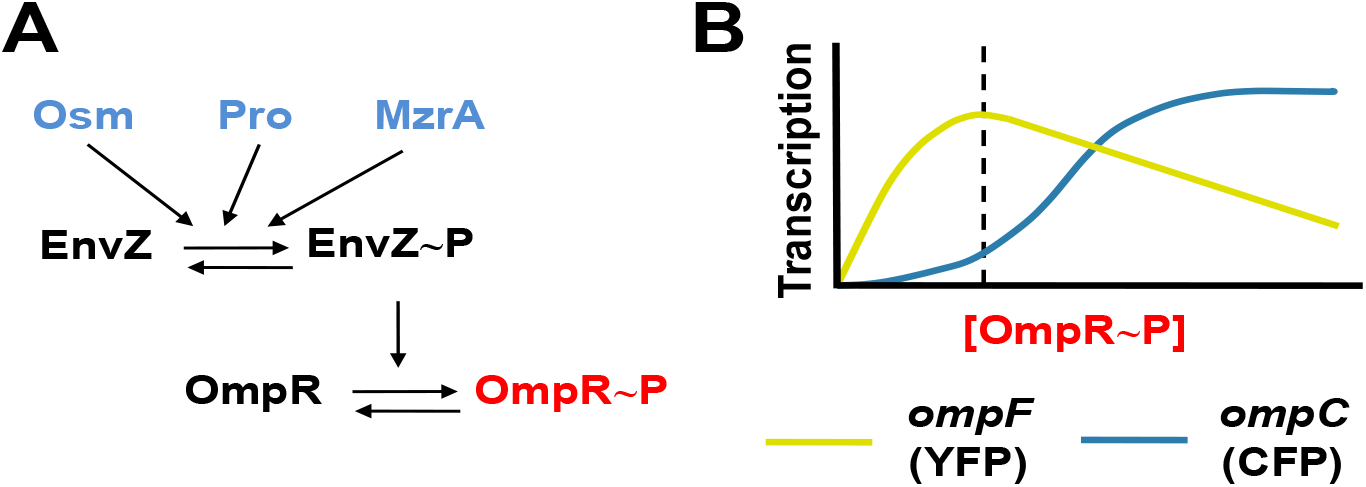
Monitoring modulation of EnvZ signal output upon stimulus perception. (A) EnvZ signal output controls porin expression. It is a bifucntional sensor histidine kinase with both kinase and phosphatase activities. The ratio of these activities is modulated by the presence of extracellular osmolarity, procaine and the absence/presence of MzrA (blue). (B) The intracellular level of phosphorylated OmpR (OmpR-P) is controlled by EnvZ signal output and OmpR-P levels govern transcription of *ompF* and *ompC*. Strains MDG147 and EPB30 (Δ*envZ)* contain transcriptional fusions of *yfp* to *ompF* (yellow) and *cfp* to *ompC* (cyan), which facilitates easy monitoring of intracellular OmpR-P levels by measuring the CFP/YFP ratio from intact cells. The dashed line indicates an estimation of the baseline OmpR-P levels from EPB30/pRD400 cells grown under the low-osmolarity regime (0% sucrose).

**Figure S2.**
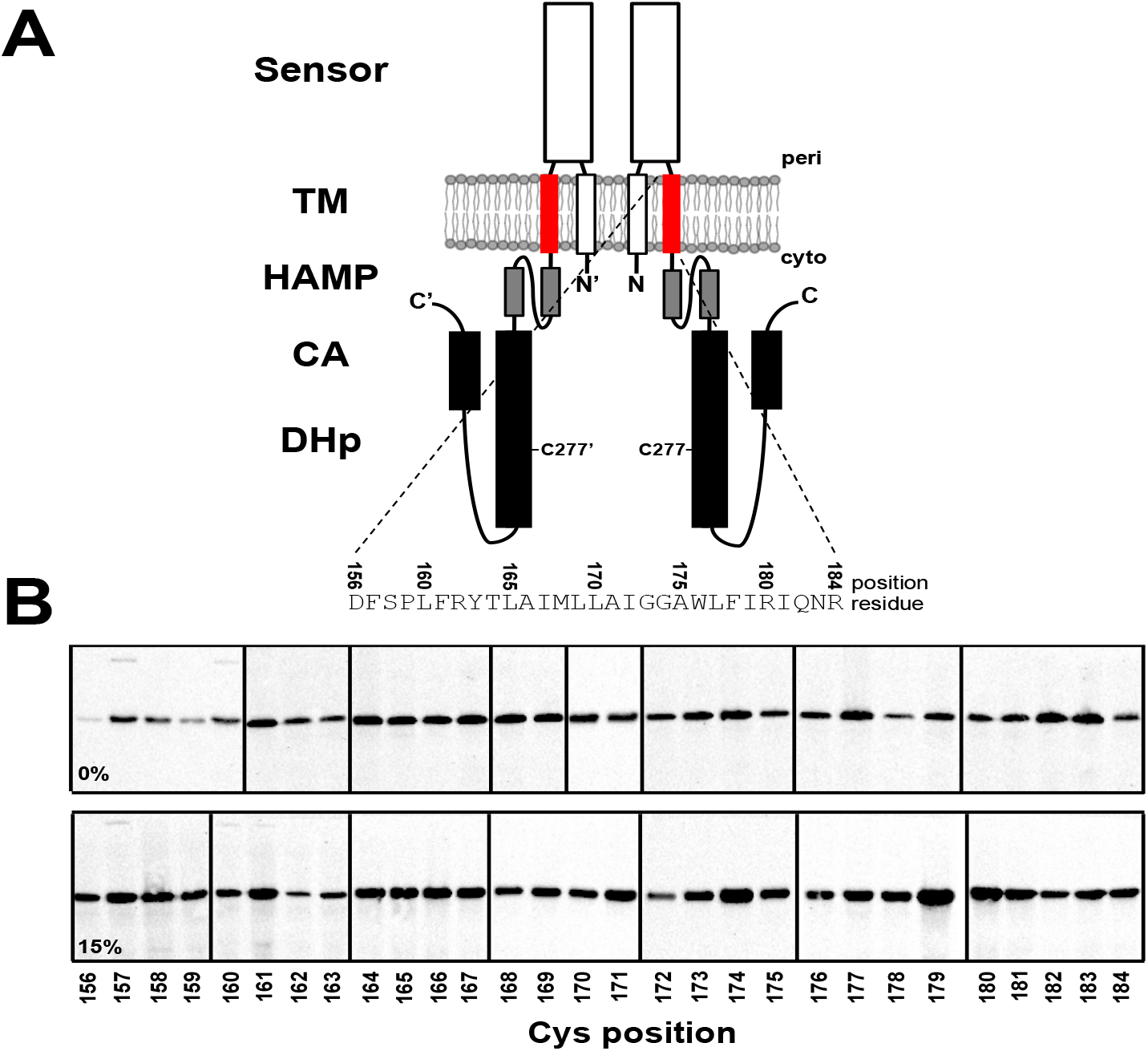
Creation of a new library of single-Cys-variants that encompasses TM2. (A) EnvZ functions as a homodimer with a cytoplasmic N-terminus, the first transmembrane helix (TM1, white), a large periplasmic domain (sensor, white), the second transmembrane helix (TM2, red), a membrane-adjacent HAMP domain (grey) and the cytoplasmic domains responsible for dimerization and histidylphosphortransfer (DHp, black) and catalytic ATPase activity (CA, black). The position of the original Cys-277 residue that was mutated to Ala to produce the Cys-less EnvZ is provided. The residues subjected to Cys substitution and their position in the primary sequence is provided. (B) Steady-state expression of EnvZ variants containing a single Cys residue within TM2. EPB30/pRD400 cells expressing one of the single-Cys-containing variants were grown under the low- (0% sucrose) or high-osmolarity (15% sucrose) regimes. Under the low-osmolarity regime, disulfide formation was observed for the F157C and L160C variants in the absence of any additional oxidizing agent. When EPB30/pRD400 cells were grown under the high-osmolarity regime, the F157C, L160C and F161C variants exhibited disulfide formation in the absence of any oxidizing agent.

**Figure S3.**
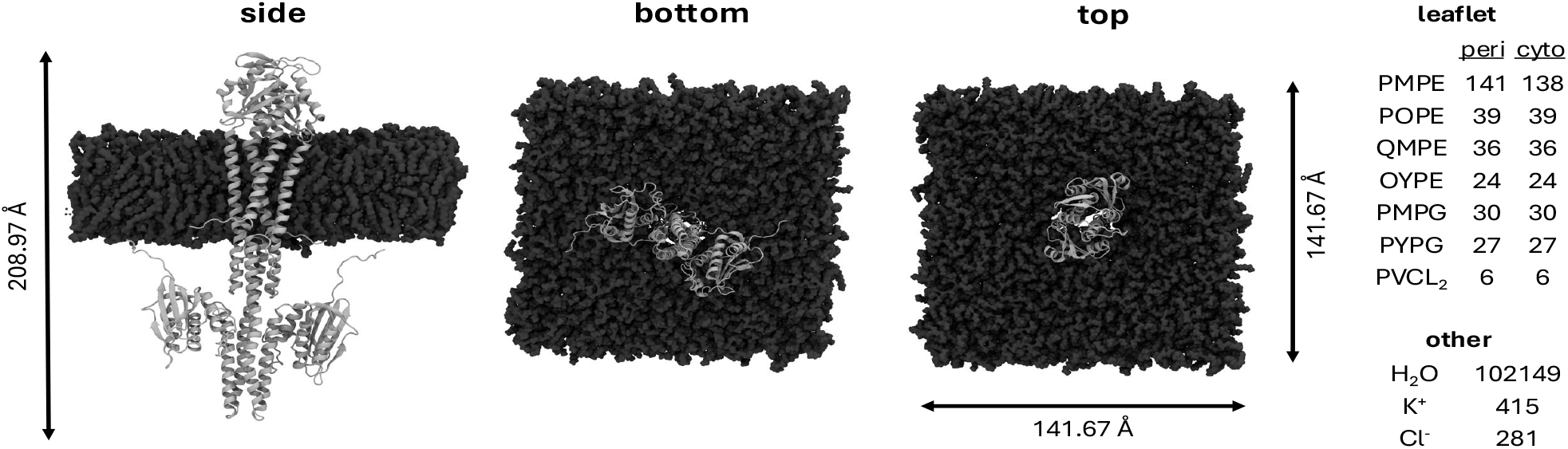
Composition and dimensions of the EnvZ_2_/G-IM system. The dimensions of the system are provided in three panels (side-view as viewed from the side, bottom-view as viewed from the cytoplasm, and top-view from the periplasm). The composition of the periplasmic and cytoplasmic leaflet and the remaining components of the system are also provided.

**Figure S4.**
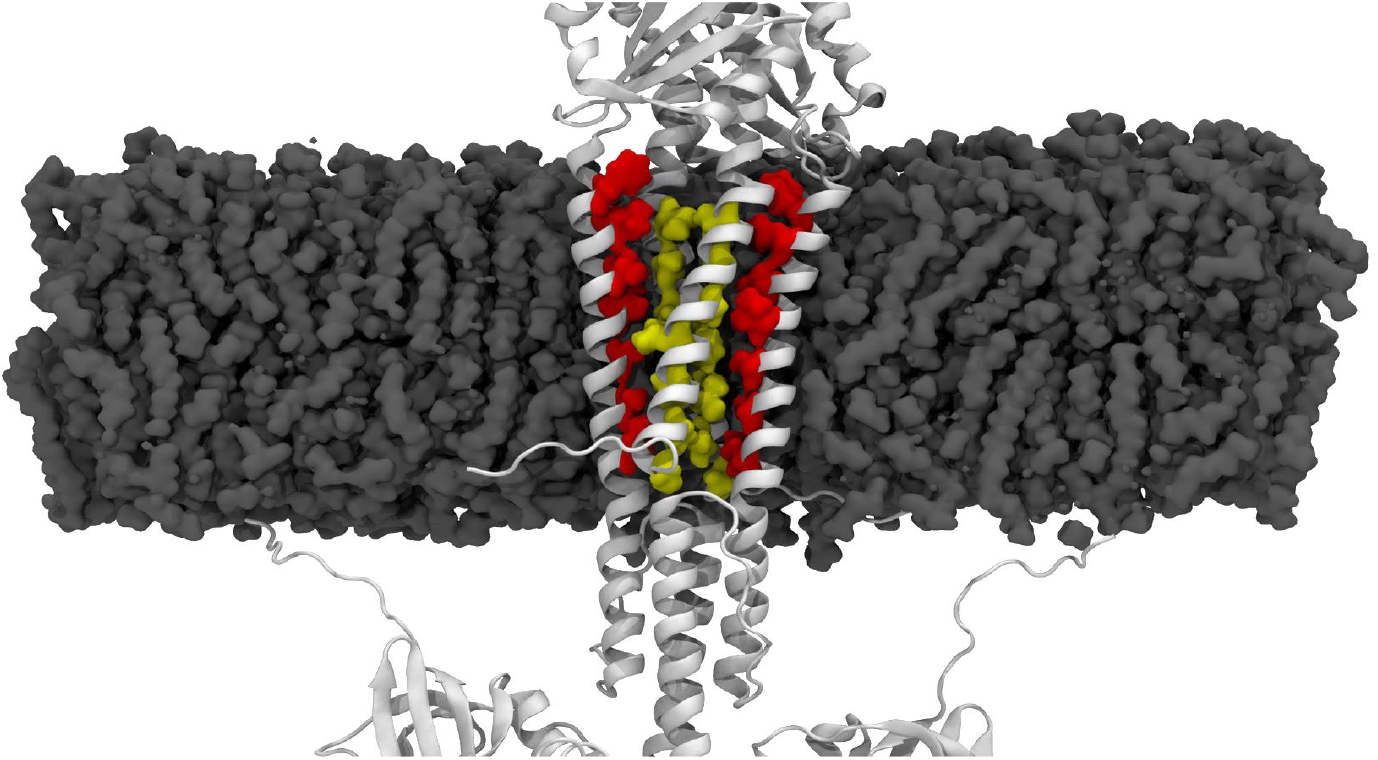
Mapping the TM1-TM1’ and the TM2-TM2’ interfaces within the EnvZ TMD. The local crosslinking maxima along TM1 and TM2, i.e. residue positions 11, 15, 19, 23, 26, 30 and 34 from TM1 and residues positions 157, 160, 164, 167, 171, 175 and 179 from TM2, are shown in yellow and red, respectively.

**Figure S5.**
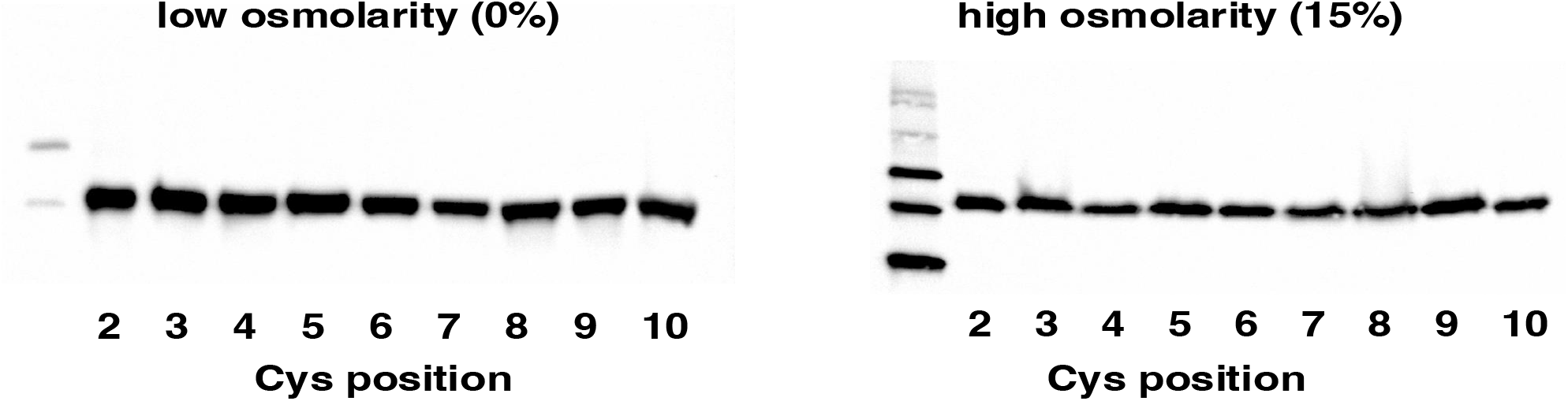
Absence of cross-linking from the cytoplasmic tail single-Cys-variants. No crosslinking is observed under the low-or high-omsolarity regimes when a Cys is present at any position between 2 and 10 in the cytoplasmic tail. Conditions for the experimentation are the same as those described in Fig. 1A.

**Figure S6.**
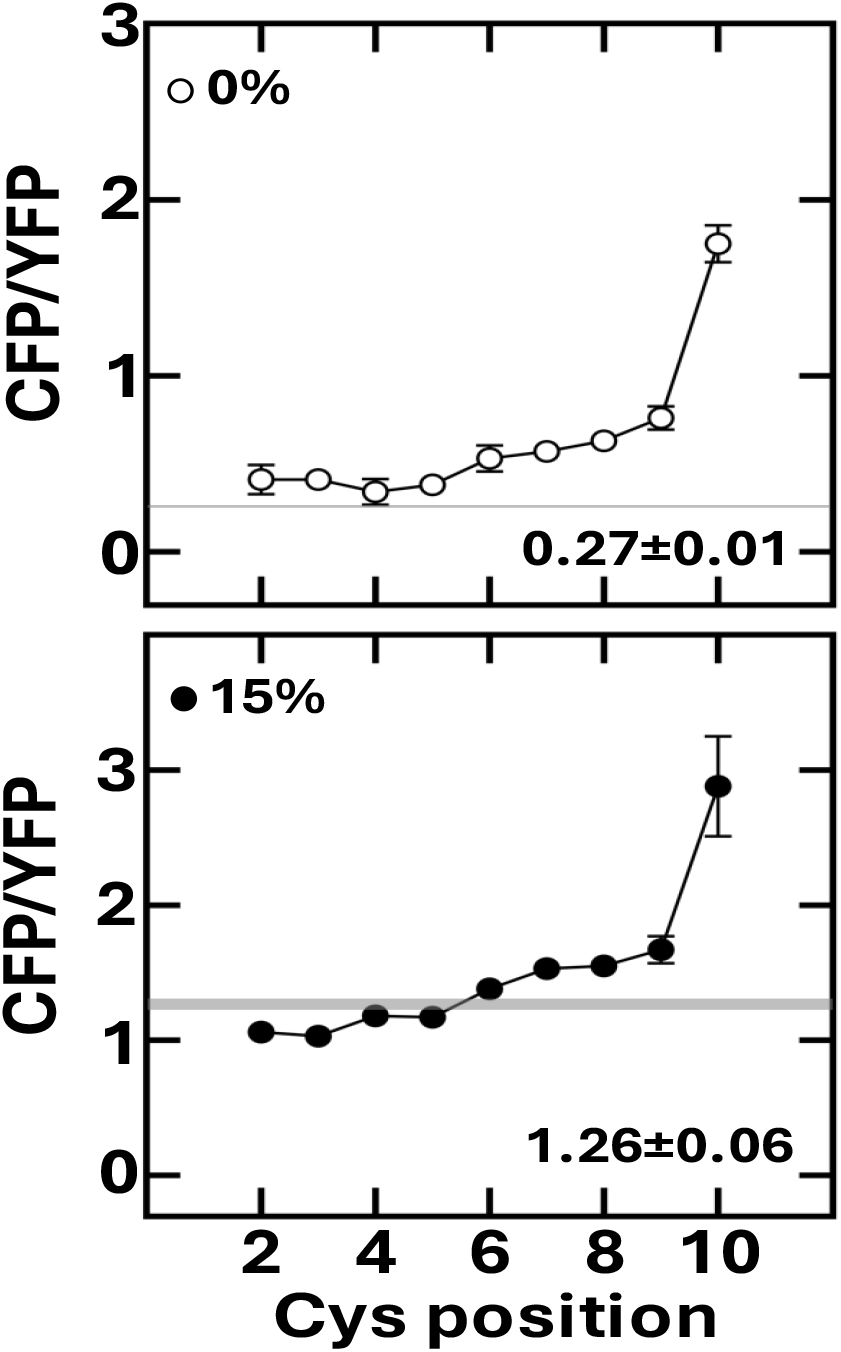
Signal output from EnvZ receptors containing a Cys residue in the cytoplasmic tail. CFP/YFP from EPB30/pRD400 cells expressing one of single-Cys variants grown under the low-osmolarity (0% sucrose, top) or high-osmolarity regime (15% sucrose, bottom). The shaded region represents signal output from the Cys-less (C277A) variant. When a Cys residue is at position 10, EnvZ signal output is significantly increased.

## References and Notes

1. M. Naghavi, S. E. Vollset, K. S. Ikuta, L. R. Swetschinski, A. P. Gray, E. E. Wool, G. Robles Aguilar, T. Mestrovic, G. Smith, C. Han, R. L. Hsu, J. Chalek, D. T. Araki, E. Chung, C. Raggi, A. Gershberg Hayoon, N. Davis Weaver, P. A. Lindstedt, A. E. Smith, U. Altay, N. V Bhattacharjee, K. Giannakis, F. Fell, B. McManigal, N. Ekapirat, J. A. Mendes, T. Runghien, O. Srimokla, A. Abdelkader, S. Abd-Elsalam, R. G. Aboagye, H. Abolhassani, H. Abualruz, U. Abubakar, H. J. Abukhadijah, S. Aburuz, A. Abu-Zaid, S. Achalapong, I. Y. Addo, V. Adekanmbi, T. E. Adeyeoluwa, Q. E. S. Adnani, L. A. Adzigbli, M. S. Afzal, S. Afzal, A. Agodi, A. J. Ahlstrom, A. Ahmad, S. Ahmad, T. Ahmad, A. Ahmadi, A. Ahmed, H. Ahmed, I. Ahmed, M. Ahmed, S. Ahmed, S. A. Ahmed, M. A. Akkaif, S. Al Awaidy, Y. Al Thaher, S. O. Alalalmeh, M. T. AlBataineh, W. A. Aldhaleei, A. A. S. Al-Gheethi, N. B. Alhaji, A. Ali, L. Ali, S. S. Ali, W. Ali, K. Allel, S. Al-Marwani, A. Alrawashdeh, A. Altaf, A. B. Al-Tammemi, J. A. Al-Tawfiq, K. H. Alzoubi, W. A. Al-Zyoud, B. Amos, J. H. Amuasi, R. Ancuceanu, J. R. Andrews, A. Anil, I. A. Anuoluwa, S. Anvari, A. E. Anyasodor, G. L. C. Apostol, J. Arabloo, M. Arafat, A. Y. Aravkin, D. Areda, A. Aremu, A. A. Artamonov, E. A. Ashley, M. O. Asika, S. S. Athari, M. M. W. Atout, T. Awoke, S. Azadnajafabad, J. M. Azam, S. Aziz, A. Y. Azzam, M. Babaei, F.-X. Babin, M. Badar, A. A. Baig, M. Bajcetic, S. Baker, M. Bardhan, H. J. Barqawi, Z. Basharat, A. Basiru, M. Bastard, S. Basu, N. S. Bayleyegn, M. A. Belete, O. O. Bello, A. Beloukas, J. A. Berkley, A. S. Bhagavathula, S. Bhaskar, S. S. Bhuyan, J. A. Bielicki, N. I. Briko, C. S. Brown, A. J. Browne, D. Buonsenso, Y. Bustanji, C. G. Carvalheiro, C.A. Castañeda-Orjuela, M. Cenderadewi, J. Chadwick, S. Chakraborty, R. M. Chandika, S. Chandy, V. Chansamouth, V. K. Chattu, A. A. Chaudhary, P. R. Ching, H. Chopra, F. R. Chowdhury, D.-T. Chu, M. Chutiyami, N. Cruz-Martins, A. G. da Silva, O. Dadras, X. Dai, S. D. Darcho, S. Das, F. P. De la Hoz, D. M. Dekker, K. Dhama, D. Diaz, B. F. R. Dickson, S. G. Djorie, M. Dodangeh, S. Dohare, K. G. Dokova, O. P. Doshi, R. K. Dowou, H. L. Dsouza, S. J. Dunachie, A. M. Dziedzic, T. Eckmanns, A. Ed-Dra, A. Eftekharimehrabad, T. C. Ekundayo, I. El Sayed, M. Elhadi, W. El-Huneidi, C. Elias, S. J. Ellis, R. Elsheikh, I. Elsohaby, C. Eltaha, B. Eshrati, M. Eslami, D. W. Eyre, A. O. Fadaka, A. F. Fagbamigbe, A. Fahim, A. Fakhri-Demeshghieh, F. O. Fasina, M. M. Fasina, A. Fatehizadeh, N. A. Feasey, A. Feizkhah, G. Fekadu, F. Fischer, I. Fitriana, K. M. Forrest, C. Fortuna Rodrigues, J. E. Fuller, M. A. Gadanya, M. Gajdács, A. P. Gandhi, E. E. Garcia-Gallo, D. O. Garrett, R. K. Gautam, M. W. Gebregergis, M. Gebrehiwot, T. G. Gebremeskel, C. Geffers, L. Georgalis, R. M. Ghazy, M. Golechha, D. Golinelli, M. Gordon, S. Gulati, R. Das Gupta, S. Gupta, V. K. Gupta, A. D. Habteyohannes, S. Haller, H. Harapan, M. L. Harrison, A. I. Hasaballah, I. Hasan, R. S. Hasan, H. Hasani, A. H. Haselbeck, M. S. Hasnain, I. I. Hassan, S. Hassan, M. S. Hassan Zadeh Tabatabaei, K. Hayat, J. He, O. E. Hegazi, M. Heidari, K. Hezam, R. Holla, M. Holm, H. Hopkins, M. M. Hossain, M. Hosseinzadeh, S. Hostiuc, N. R. Hussein, L. D. Huy, E.D. Ibáñez-Prada, A. Ikiroma, I. M. Ilic, S. M. S. Islam, F. Ismail, N. E. Ismail, C. D. Iwu, C. J. Iwu-Jaja, A. Jafarzadeh, F. Jaiteh, R. Jalilzadeh Yengejeh, R. D. G. Jamora, J. Javidnia, T. Jawaid, A. W. J. Jenney, H. J. Jeon, M. Jokar, N. Jomehzadeh, T. Joo, N. Joseph, Z. Kamal, K. K. Kanmodi, R. S. Kantar, J. A. Kapisi, I. M. Karaye, Y. S. Khader, H. Khajuria, N. Khalid, F. Khamesipour, A. Khan, M. J. Khan, M. T. Khan, V. Khanal, F. F. Khidri, J. Khubchandani, S. Khusuwan, M. S. Kim, A. Kisa, V. A. Korshunov, F. Krapp, R. Krumkamp, M. Kuddus, M. Kulimbet, D. Kumar, E. A. P. Kumaran, A. Kuttikkattu, H. H. Kyu, I. Landires, B. K. Lawal, T. T. T. Le, I. M. Lederer, M. Lee, S. W. Lee, A. Lepape, T. L. Lerango, V. S. Ligade, C. Lim, S. S. Lim, L. W. Limenh, C. Liu, X. Liu, X. Liu, M. J. Loftus, H. I. M Amin, K. L. Maass, S. B. Maharaj, M. A. Mahmoud, P. Maikanti-Charalampous, O. M. Makram, K. Malhotra, A. A. Malik, G. D. Mandilara, F. Marks, B. A. Martinez-Guerra, M. Martorell, H. Masoumi-Asl, A. G. Mathioudakis, J. May, T. A. McHugh, J. Meiring, H. N. Meles, A. Melese, E. B. Melese, G. Minervini, N. S. Mohamed, S. Mohammed, S. Mohan, A. H. Mokdad, L. Monasta, A. Moodi Ghalibaf, C. E. Moore, Y. Moradi, E. Mossialos, V. Mougin, G. D. Mukoro, F. Mulita, B. Muller-Pebody, E. Murillo-Zamora, S. Musa, P. Musicha, L. A. Musila, S. Muthupandian, A. J. Nagarajan, P. Naghavi, F. Nainu, T. S. Nair, H. H. R. Najmuldeen, Z. S. Natto, J. Nauman, B. P. Nayak, G. T. Nchanji, P. Ndishimye, I. Negoi, R. I. Negoi, S. A. Nejadghaderi, Q. P. Nguyen, E. A. Noman, D. C. Nwakanma, S. O’Brien, T. J. Ochoa, I. A. Odetokun, O. A. Ogundijo, T. R. Ojo-Akosile, S. R. Okeke, O. C. Okonji, A. T. Olagunju, A. Olivas-Martinez, A. A. Olorukooba, P. Olwoch, K. I. Onyedibe, E. Ortiz-Brizuela, O. Osuolale, P. Ounchanum, O. T. Oyeyemi, M. P. P A, J. L. Paredes, R. R. Parikh, J. Patel, S. Patil, S. Pawar, A. Y. Peleg, P. Peprah, J. Perdigão, C. Perrone, I.-R. Petcu, K. Phommasone, Z. Z. Piracha, D. Poddighe, A. J. Pollard, R. Poluru, A. Ponce-De-Leon, J. Puvvula, F. N. Qamar, N. H. Qasim, C. D. Rafai, P. Raghav, L. Rahbarnia, F. Rahim, V. Rahimi-Movaghar, M. Rahman, M. A. Rahman, H. Ramadan, S. K. Ramasamy, P. S. Ramesh, P. W. Ramteke, R. K. Rana, U. Rani, M.-M. Rashidi, D. Rathish, S. Rattanavong, S. Rawaf, E. M. M. Redwan, L. F. Reyes, T. Roberts, J. V Robotham, V. D. Rosenthal, A. G. Ross, N. Roy, K. E. Rudd, C. J. Sabet, B. A. Saddik, M. R. Saeb, U. Saeed, S. Saeedi Moghaddam, W. Saengchan, M. Safaei, A. Saghazadeh, N. Saheb Sharif-Askari, A. Sahebkar, S. S. Sahoo, M. Sahu, M. Saki, N. Salam, Z. Saleem, M. A. Saleh, Y. L. Samodra, A. M. Samy, A. Saravanan, M. Satpathy, A. E. Schumacher, M. Sedighi, S. Seekaew, M. Shafie, P. A. Shah, S. Shahid, M. J. Shahwan, S. Shakoor, N. Shalev, M. A. Shamim, M. A. Shamshirgaran, A. Shamsi, A. Sharifan, R. P. Shastry, M. Shetty, A. Shittu, S. Shrestha, E. E. Siddig, T. Sideroglou, J. Sifuentes-Osornio, L. M. L. R. Silva, E. A. F. Simões, A. J. H. Simpson, A. Singh, S. Singh, R. Sinto, S. S. M. Soliman, S. Soraneh, N. Stoesser, T. Z. Stoeva, C. K. Swain, L. Szarpak, S. S. T Y S. Tabatabai, C. Tabche, Z. M.-A. Taha, K.-K. Tan, N. Tasak, N. Y. Tat, A. Thaiprakong, P. Thangaraju, C. C. Tigoi, K. Tiwari, M. R. Tovani-Palone, T. H. Tran, M. Tumurkhuu, P. Turner, A. J. Udoakang, A. Udoh, N. Ullah, S. Ullah, A. G. Vaithinathan, M. Valenti, T. Vos, H. T. L. Vu, Y. Waheed, A. S. Walker, J. L. Walson, T. Wangrangsimakul, K. G. Weerakoon, H. F. L. Wertheim, P. C. M. Williams, A. A. Wolde, T. M. Wozniak, F. Wu, Z. Wu, M. K. K. Yadav, S. Yaghoubi, Z. S. Yahaya, A. Yarahmadi, S. Yezli, Y. E. Yismaw, D. K. Yon, C.-W. Yuan, H. Yusuf, F. Zakham, G. Zamagni, H. Zhang, Z.-J. Zhang, M. Zielińska, A. Zumla, S. H. H. Zyoud, S. H. Zyoud, S. I. Hay, A. Stergachis, B. Sartorius, B. S. Cooper, C. Dolecek, C. J. L. Murray, Global burden of bacterial antimicrobial resistance 1990-2021: a systematic analysis with forecasts to 2050. The Lancet 404, 1199–1226 (2024).

2. J. O’Neill, “Antimicrobial Resistance: Tackling a Crisis for the Health and Wealth of Nations.” (2014).

3. N. Hiroshi, Molecular Basis of Bacterial Outer Membrane Permeability Revisited. Microbiology and Molecular Biology Reviews 67, 593–656 (2003).

4. H. Nikaido, Prevention of Drug Access to Bacterial Targets: Permeability Barriers and Active Efflux. Science (1979) 264, 382–388 (1994).

5. A. H. Delcour, Solute uptake through general porins. FBL 8, 1055–1071 (2003).

6. G. E. Schulz, The structure of bacterial outer membrane proteins. Biochimica et Biophysica Acta (BBA) - Biomembranes 1565, 308–317 (2002).

7. L. A. Egger, M. Inouye, Purification and Characterization of the Periplasmic Domain of EnvZ Osmosensor inEscherichia coli. Biochem Biophys Res Commun 231, 68–72 (1997).

8. S. A. Forst, D. L. Roberts, Signal transduction by the EnvZ-OmpR phosphotransfer system in bacteria. Res Microbiol 145, 363–373 (1994).

9. H. Gerken, E. S. Charlson, E. M. Cicirelli, L. J. Kenney, R. Misra, MzrA: a novel modulator of the EnvZ/OmpR two-component regulon. Mol Microbiol 72, 1408–1422 (2009).

10. T. Mizuno, His-Asp Phosphotransfer Signal Transduction1. The Journal of Biochemistry 123, 555–563 (1998).

11. G. Henri, M. Rajeev, MzrA-EnvZ Interactions in the Periplasm Influence the EnvZ/OmpR Two-Component Regulon. J Bacteriol 192, 6271–6278 (2010).

12. S. A. Forst, J. Delgado, M. Inouye, DNA-binding properties of the transcription activator (OmpR) for the upstream sequences of ompF in Escherichia coli are altered by envZ mutations and medium osmolarity. J Bacteriol 171, 2949–2955 (1989).

13. L. Chung-Yu, I. M. M, Differential Expression of the OmpF and OmpC Porin Proteins in Escherichia coli K-12 Depends upon the Level of Active OmpR. J Bacteriol 180, 171–174 (1998).

14. F. D. Russo, T. J. Silhavy, EnvZ controls the concentration of phosphorylated OmpR to mediate osmoregulation of the porin genes. J Mol Biol 222, 567–580 (1991).

15. S. Hernández-Allés, M. del C. Conejo, A. Pascual, J. M. Tomás, V. J. Benedí, L. Martínez-Martínez, Relationship between outer membrane alterations and susceptibility to antimicrobial agents in isogenic strains of Klebsiella pneumoniae. Journal of Antimicrobial Chemotherapy 46, 273–277 (2000).

16. J.G. A M. D. M C. Nancy, Role of β-Lactamases and Porins in Resistance to Ertapenem and Other β-Lactams in Klebsiella pneumoniae. Antimicrob Agents Chemother 48, 3203–3206 (2004).

17. K. F. M D.-H. Fadia, S. Wenchi, G. T. D, High-Level Carbapenem Resistance in a Klebsiella pneumoniae Clinical Isolate Is Due to the Combination of blaACT-1 β-Lactamase Production, Porin OmpK35/36 Insertional Inactivation, and Down-Regulation of the Phosphate Transport Porin PhoE. Antimicrob Agents Chemother 50, 3396–3406 (2006).

18. M. Ana, P. Virginia, G. Laura, H. Olga, A. J. Ignacio, A. Sebastián, B. Nuria, P. J. L O. Antonio, Characterization of a Large Outbreak by CTX-M-1-Producing Klebsiella pneumoniae and Mechanisms Leading to In Vivo Carbapenem Resistance Development. J Clin Microbiol 44, 2831–2837 (2006).

19. L. Martínez-Martínez, M. C. Conejo, A. Pascual, S. Hernández-Allés, S. Ballesta, R. de A.-R. E. V. J. Benedí, E. J. Perea, Activities of Imipenem and Cephalosporins against Clonally Related Strains of Escherichia coli Hyperproducing Chromosomal β-Lactamase and Showing Altered Porin Profiles. Antimicrob Agents Chemother 44, 2534–2536 (2000).

20. A. Loli, L. S. Tzouvelekis, E. Tzelepi, A. Carattoli, A. C. Vatopoulos, P. T. Tassios, V. Miriagou, Sources of diversity of carbapenem resistance levels in Klebsiella pneumoniae carrying blaVIM-1. Journal of Antimicrobial Chemotherapy 58, 669–672 (2006).

21. E. Elliott, A. J. Brink, J. van Greune, Z. Els, N. Woodford, J. Turton, M. Warner, D. M. Livermore, In Vivo Development of Ertapenem Resistance in a Patient with Pneumonia Caused by Klebsiella pneumoniae with an Extended-Spectrum β-Lactamase. Clinical Infectious Diseases 42, e95–e98 (2006).

22. M. Adler, M. Anjum, D. I. Andersson, L. Sandegren, Combinations of mutations in envZ, ftsI, mrdA, acrB and acrR can cause high-level carbapenem resistance in Escherichia coli. Journal of Antimicrobial Chemotherapy 71, 1188–1198 (2016).

23. L. C. Wang, L. K. Morgan, P. Godakumbura, L. J. Kenney, G. S. Anand, The inner membrane histidine kinase EnvZ senses osmolality via helix-coil transitions in the cytoplasm. EMBO J 31, 2648-2659–2659 (2012).

24. A. Heininger, R. Yusuf, R. J. Lawrence, R. R. Draheim, Identification of transmembrane helix 1 (TM1) surfaces important for EnvZ dimerisation and signal output. Biochimica et Biophysica Acta (BBA) - Biomembranes 1858, 1868–1875 (2016).

25. T. Hessa, N. M. Meindl-Beinker, A. Bernsel, H. Kim, Y. Sato, M. Lerch-Bader, I. Nilsson, S. H. White, G. von Heijne, Molecular code for transmembrane-helix recognition by the Sec61 translocon. Nature 450, 1026–1030 (2007).

26. A. Krogh, B. Larsson, G. von Heijne, E. L. L. Sonnhammer, Predicting transmembrane protein topology with a hidden markov model: application to complete genomes11Edited by F. Cohen. J Mol Biol 305, 567– 580 (2001).

27. J. Jumper, R. Evans, A. Pritzel, T. Green, M. Figurnov, O. Ronneberger, K. Tunyasuvunakool, R. Bates, A. Žídek, A. Potapenko, A. Bridgland, C. Meyer, S. A. A. Kohl, A. J. Ballard, A. Cowie, B. Romera-Paredes, S. Nikolov, R. Jain, J. Adler, T. Back, S. Petersen, D. Reiman, E. Clancy, M. Zielinski, M. Steinegger, M. Pacholska, T. Berghammer, S. Bodenstein, D. Silver, O. Vinyals, A. W. Senior, K. Kavukcuoglu, P. Kohli, D. Hassabis, Highly accurate protein structure prediction with AlphaFold. Nature 596, 583–589 (2021).

28. R. Evans, M. O’Neill, A. Pritzel, N. Antropova, A. Senior, T. Green, A. Žídek, R. Bates, S. Blackwell, J. Yim, O. Ronneberger, S. Bodenstein, M. Zielinski, A. Bridgland, A. Potapenko, A. Cowie, K. Tunyasuvunakool, R. Jain, E. Clancy, P. Kohli, J. Jumper, D. Hassabis, Protein complex prediction with AlphaFold-Multimer. bioRxiv, 2021.10.04.463034 (2021).

29. I. D. Pogozheva, G. A. Armstrong, L. Kong, T. J. Hartnagel, C. A. Carpino, S. E. Gee, D. M. Picarello, A. S. Rubin, J. Lee, S. Park, A. L. Lomize, W. Im, Comparative Molecular Dynamics Simulation Studies of Realistic Eukaryotic, Prokaryotic, and Archaeal Membranes. J Chem Inf Model 62, 1036–1051 (2022).

30. S. Jo, T. Kim, V. G. Iyer, W. Im, CHARMM-GUI: A web-based graphical user interface for CHARMM. J Comput Chem 29, 1859–1865 (2008).

31. J. Lee, X. Cheng, J. M. Swails, M. S. Yeom, P. K. Eastman, J. A. Lemkul, S. Wei, J. Buckner, J. C. Jeong, Y. Qi, S. Jo, V. S. Pande, D. A. Case, C. L. I. I. I. Brooks, A. D. Jr. MacKerell, J. B. Klauda, W. Im, CHARMM-GUI Input Generator for NAMD, GROMACS, AMBER, OpenMM, and CHARMM/OpenMM Simulations Using the CHARMM36 Additive Force Field. J Chem Theory Comput 12, 405–413 (2016).

32. B. R. Brooks, C. L. Brooks III, A. D. Mackerell Jr., L. Nilsson, R. J. Petrella, B. Roux, Y. Won, G. Archontis, C. Bartels, S. Boresch, A. Caflisch, L. Caves, Q. Cui, A. R. Dinner, M. Feig, S. Fischer, J. Gao, M. Hodoscek, W. Im, K. Kuczera, T. Lazaridis, J. Ma, V. Ovchinnikov, E. Paci, R. W. Pastor, C. B. Post, J.Z. Pu, M. Schaefer, B. Tidor, R. M. Venable, H. L. Woodcock, X. Wu, W. Yang, D. M. York, M. Karplus, CHARMM: The biomolecular simulation program. J Comput Chem 30, 1545–1614 (2009).

33. E. L. Wu, X. Cheng, S. Jo, H. Rui, K. C. Song, E. M. Dávila-Contreras, Y. Qi, J. Lee, V. Monje-Galvan, R. M. Venable, J. B. Klauda, W. Im, CHARMM-GUI Membrane Builder toward realistic biological membrane simulations. J Comput Chem 35, 1997–2004 (2014).

34. S. Jo, T. Kim, W. Im, Automated Builder and Database of Protein/Membrane Complexes for Molecular Dynamics Simulations. PLoS One 2, e880.(2007).

35. S. Feng, S. Park, Y. K. Choi, W. Im, CHARMM-GUI Membrane Builder: Past, Current, and Future Developments and Applications. J Chem Theory Comput 19, 2161–2185 (2023).

36. J. C. Phillips, D. J. Hardy, J. D. C. Maia, J. E. Stone, J. V Ribeiro, R. C. Bernardi, R. Buch, G. Fiorin, J. Hénin, W. Jiang, R. McGreevy, M. C. R. Melo, B. K. Radak, R. D. Skeel, A. Singharoy, Y. Wang, B. Roux, A. Aksimentiev, Z. Luthey-Schulten, L.V Kalé, K. Schulten, C. Chipot, E. Tajkhorshid, Scalable molecular dynamics on CPU and GPU architectures with NAMD. J Chem Phys 153, 044130 (2020).

37. S. D. Goldberg, G. D. Clinthorne, M. Goulian, W. F. DeGrado, Transmembrane polar interactions are required for signaling in the Escherichia coli sensor kinase PhoQ. Proceedings of the National Academy of Sciences 107, 8141–8146 (2010).

38. V. I. Gordeliy, J. Labahn, R. Moukhametzianov, R. Efremov, J. Granzin, R. Schlesinger, G. Büldt, T. Savopol, A. J. Scheidig, J. P. Klare, M. Engelhard, Molecular basis of transmembrane signalling by sensory rhodopsin II–transducer complex. Nature 419, 484–487 (2002).

39. T. Lemmin, C. S. Soto, G. Clinthorne, W. F. DeGrado, M. Dal Peraro, Assembly of the Transmembrane Domain of E. coli PhoQ Histidine Kinase: Implications for Signal Transduction from Molecular Simulations. PLoS Comput Biol 9, e1002878. (2013).

40. I. Gushchin, I. Melnikov, V. Polovinkin, A. Ishchenko, A. Yuzhakova, P. Buslaev, G. Bourenkov, S. Grudinin, E. Round, T. Balandin, V. Borshchevskiy, D. Willbold, G. Leonard, G. Büldt, A. Popov, V. Gordeliy, Mechanism of transmembrane signaling by sensor histidine kinases. Science (1979) 356, eaah6345 (2017).

41. S. Lazaridi, J. Yuan, T. Lemmin, Atomic insights into the signaling landscape of <em>E. coli</em> PhoQ Histidine Kinase from Molecular Dynamics simulations. bioRxiv, 2024.04.19.590235 (2024).

42. R. Yusuf, R. R. Draheim, Employing aromatic tuning to modulate output from two-component signaling circuits. J Biol Eng 9, 7 (2015).

43. M. H. H. Nørholm, G. von Heijne, R. R. Draheim, Forcing the Issue: Aromatic Tuning Facilitates Stimulus-Independent Modulation of a Two-Component Signaling Circuit. ACS Synth Biol 4, 474–481 (2015).

44. S. C. Botelho, K. Enquist, G. von Heijne, R. R. Draheim, Differential repositioning of the second transmembrane helices from E. coli Tar and EnvZ upon moving the flanking aromatic residues. Biochimica et Biophysica Acta (BBA) - Biomembranes 1848, 615–621 (2015).

45. C. A. Adase, R. R. Draheim, M. D. Manson, The Residue Composition of the Aromatic Anchor of the Second Transmembrane Helix Determines the Signaling Properties of the Aspartate/Maltose Chemoreceptor Tar of Escherichia coli. Biochemistry 51, 1925–1932 (2012).

46. R. R. Draheim, A. F. Bormans, R. Lai, M. D. Manson, Tryptophan Residues Flanking the Second Transmembrane Helix (TM2) Set the Signaling State of the Tar Chemoreceptor. Biochemistry 44, 1268– 1277 (2005).

47. C. A. Adase, R. R. Draheim, G. Rueda, R. Desai, M. D. Manson, Residues at the Cytoplasmic End of Transmembrane Helix 2 Determine the Signal Output of the TarEc Chemoreceptor. Biochemistry 52, 2729– 2738 (2013).

48. R. R. Draheim, A. F. Bormans, R.-Z. Lai, M. D. Manson, Tuning a Bacterial Chemoreceptor with Protein−Membrane Interactions. Biochemistry 45, 14655–14664 (2006).

49. R. Yusuf, R. J. Lawrence, L. V Eke, R. R. Draheim, “Tuning Chemoreceptor Signaling by Positioning Aromatic Residues at the Lipid–Aqueous Interface” in Bacterial Chemosensing: Methods and Protocols, M.D. Manson, Ed. (Springer New York, New York, NY, 2018), pp. 147–158.

50. M. J. Casadaban, S. N. Cohen, Analysis of gene control signals by DNA fusion and cloning in Escherichia coli. J Mol Biol 138, 179–207 (1980).

51. M. S. Guyer, R. R. Reed, J. A. Steitz, K. B. Low, Identification of a Sex-factor-affinity Site in E. coli as γδ. Cold Spring Harb Symp Quant Biol 45, 135–140 (1981).

52. B. Eric, S. T. J G. Mark, Continuous Control in Bacterial Regulatory Circuits. J Bacteriol 186, 7618–7625 (2004).

53. A. Siryaporn, M. Goulian, Cross-talk suppression between the CpxA–CpxR and EnvZ–OmpR two-component systems in E. coli. Mol Microbiol 70, 494–506 (2008).

54. R.-Z. Lai, A. F. Bormans, R. R. Draheim, G. A. Wright, M. D. Manson, The Region Preceding the C-Terminal NWETF Pentapeptide Modulates Baseline Activity and Aspartate Inhibition of Escherichia coli Tar. Biochemistry 47, 13287–13295 (2008).

55. E. Batchelor, M. Goulian,Robustness and the cycle of phosphorylation and dephosphorylation in a two-component regulatory system. Proceedings of the National Academy of Sciences 100, 691–696 (2003).

56. W. Hsing, T. J. Silhavy, Function of conserved histidine-243 in phosphatase activity of EnvZ, the sensor for porin osmoregulation in Escherichia coli. J Bacteriol 179, 3729–3735 (1997).

57. C. B. J, D. R. R, W. R. B, N. Cameran, S. R. C. M. M. D, CheZ Phosphatase Localizes to Chemoreceptor Patches via CheA-Short. J Bacteriol 185, 2354–2361 (2003).

58. J. A. Southern, D. F. Young, F. Heaney, W. K. Baumgärtner, R. E. Randall, Identification of an epitope on the P and V proteins of simian virus 5 that distinguishes between two isolates with different biological characteristics. Journal of General Virology 72, 1551–1557 (1991).

59. J. Miller, A Short Course in Bacterial Genetics: A Laboratory Manual and Handbook for Escherichia Coli and Related Bacteria. (Cold Spring Harbor Laboratory Press, Plainview, NY, 1992).

60. F. Ausubel, R. Brent, R. Kingston, D. Moore, J. Seidman, J. Smith, K. Struhl, Current Protocols in Molecular Biology (Wiley, New York, NY, 1998).

61. C. A. Schneider, W. S. Rasband, K. W. Eliceiri, NIH Image to ImageJ: 25 years of image analysis. Nat Methods 9, 671–675 (2012).

62. P. J. Steinbach, B. R. Brooks, New spherical-cutoff methods for long-range forces in macromolecular simulation. J Comput Chem 15, 667–683 (1994).

63. U. Essmann, L. Perera, M. L. Berkowitz, T. Darden, H. Lee, L. G. Pedersen, A smooth particle mesh Ewald method. J Chem Phys 103, 8577–8593 (1995).

64. K.-H. Chow, D. M. Ferguson, Isothermal-isobaric molecular dynamics simulations with Monte Carlo volume sampling. Comput Phys Commun 91, 283–289 (1995).

65. N. Michaud-Agrawal, E. J. Denning, T. B. Woolf, O. Beckstein, MDAnalysis: A toolkit for the analysis of molecular dynamics simulations. J Comput Chem 32, 2319–2327 (2011).

